# Trajectory changes are susceptible to change blindness manipulations

**DOI:** 10.1101/391359

**Authors:** Matt Jaquiery, Nora Andermane, Ron Chrisley

## Abstract

People routinely fail to notice that things have changed in a visual scene if they do not perceive the changes in the process of occurring, a phenomenon known as ‘change blindness’ (1,2). The majority of lab-based change blindness studies use static stimuli and require participants to identify simple changes such as alterations in stimulus orientation or scene composition. This study uses a ‘flicker’ paradigm adapted for dynamic stimuli which allowed for both simple orientation changes and more complex trajectory changes. Participants were required to identify a moving rectangle which underwent one of these changes against a background of moving rectangles which did not. The results demonstrated that participants’ ability to correctly identify the target deteriorated with the presence of a visual mask and a larger number of distractor objects, consistent with findings in previous change blindness work.

The study provides evidence that the flicker paradigm can be used to induce change blindness with dynamic stimuli, and that changes to predictable trajectories are detected or missed in the similar way as orientation changes.

## Introduction

People routinely fail to notice that objects have changed in a visual scene if they do not perceive the changes in the process of occurring, a phenomenon known as ‘change blindness’ (1,2). The majority of lab-based change blindness studies use static stimuli and require participants to identify simple changes such as alterations in stimulus orientation or scene composition (3–5), though others use more complex and realistic environments, especially driving simulators (6–8). This study examines whether changes to dynamic properties are detected or missed in the same way as changes to static properties.

Changes to static properties (e.g. the presence of a stimulus, or its orientation) are most readily detected when the transients (moment-to-moment variations) accompanying a change prompt an explicit comparison between a stored representation of a stimulus and its current presentation (9–11). Change blindness frequently occurs when this process is disrupted. If the representation of a stimulus includes information on dynamic properties (e.g. the trajectory along which a stimulus is travelling), change blindness would be expected to occur for changes to dynamic as well as static stimulus properties. Common methodologies for inducing change blindness prevent transient registration, typically by masking (12) or eliminating transients (1,13,14). In addition to masking transients, exhaustion of working memory capacity is required to produce change blindness effects reliably (15), with the contents of working memory exhibiting resistance to change blindness (2,16), a phenomenon which is stable enough to allow change blindness task performance to act as a guide to working memory contents in attentional bias studies (17–19).

Change blindness as deployed in the study of other phenomena (e.g. attentional biases) is reasonably well understood, but a gap exists between these structured laboratory experiments and the more sophisticated simulator-based and natural-world experiments (1,20,21). The use of dynamic paradigms such as video footage (22,23) or programmed displays (24) is required for a detailed examination of the processes underpinning change blindness within busy, continually changing visual fields like those typical of everyday life (25).

Two key areas of enquiry are addressable with the use of dynamic stimuli: the nature of competition between transients in exogenous orienting (‘grabbing’) of attention; and the existence of an ability to make discriminations between patterns of transients. The first of these areas is to some extent already established by the existence of change blindness paradigms in which attention is directed away from transients by the application of ‘mudsplashes’ or similar distractors (12): the more prominent transients accompanying the mudsplashes outcompete those accompanying the change in the target stimulus leading to observable change blindness. The second area of enquiry, discrimination between patterns of transients, is only available in dynamic scenes where changes are already occurring, and requires the detection not of specific transients but of a change in the pattern of those transients. A change in a pattern of transients signifies a change in the way in which a change is occurring (e.g. an acceleration or a change in the direction of movement).

Change blindness to changes in trajectory has been demonstrated in macaque monkeys in a study which used a change in the flow direction of dots within a field, with distractor fields in which the flow direction remained constant also visible (26). The present experiment establishes a related finding in humans, namely that trajectory changes can drive attentional mechanisms in the same manner as orientation changes, demonstrating detection of and blindness to changes in an object’s dynamic properties where the alterations to the patterns of transients are detected or undetected, respectively.

## Materials & Methods

### Study structure

The results presented below comprise an online study and a lab-based replication. Both studies used the same materials and methods; however, the experimental environment and presentation was standardized for the lab cohort.

### Participants

Participants (*N* online = 42, *N* lab = 16) were recruited to the study. Online participants were recruited via university mailing lists and social networking websites, while lab participants were recruited via the mailing lists and word-of-mouth. Prior to beginning the task participants were informed that no personally identifying information would be recorded, that participation was voluntary and could be halted at any time, and that they would be identified by means of temporary browser cookies. The experiment was administered online and demographic data such as age, education, and gender identity were not collected. It is likely the majority of the online cohort were undergraduates. The lab cohort were undergraduate and postgraduate students at the University of Sussex.

Target sample size for the online cohort was determined by precedent. Similar change blindness paradigms administered under laboratory conditions have used sample sizes in the 10-20 range (1,2,5,12,17,18,27). Given the reduction in precision accompanying the novel online administration in the present study, the previous range was doubled, resulting in a target sample size of 20-40. Active recruitment lasted two weeks, although participants were free to enter the study until the pre-established one-month data collection window had closed. Power analysis was used to determine the number of participants required for the lab-based study to produce 95% power for detecting the key interaction between masking and load as demonstrated in the online data.

### Ethics

This research was conducted in accordance with the ethics procedures of the University of Sussex School of Life Sciences, and approved by the University of Sussex Life Sciences School Research Ethics Officer on behalf of the Cluster-based Research Ethics Committee Ethical Review Application ID (ER/MJ261/1). After being briefed about the content and nature of the task the participants were required to signal consent by following a hyperlink to the task page. Participants were informed that participation was voluntary and that they were free to stop at any time. Participation was unpaid, but the lab cohort were entitled to receive a small amount of course credit.

### Task

The participants completed the task by visiting a webpage which used JavaScript to deliver a dynamic version of a ‘flicker’ paradigm (2). In a typical flicker paradigm, as in most change blindness paradigms, the task is to detect changes between two stimuli. In the flicker paradigm the participant is shown one stimulus, then the other, repeatedly. Importantly, the stimuli presentations are separated by a brief presentation of a blank screen (the ‘mask’). If the stimuli are switched without a mask the parts of the scene that are different produce visual transients, attracting attention to the location of the change. If, however, the switch is accompanied by the mask, the offset of the first stimulus and the onset of the second both produce transients throughout the visual scene, resulting in no net increase in attention to the location of the change.

The task page presented participants with a 700×700 pixel working area. The initial stimulus consisted of a number of 50×25 pixel rectangles with randomly selected colours moved at 150 pixels/second along a straight-line trajectory (Fig 1a). On low load trials there were 2 rectangles; on high load trials there were 6. The direction of movement for each rectangle was determined randomly, subject to the constraints that: a) the rectangle could travel along the selected trajectory for the duration of the trial without leaving the working area; and b) there existed at least one possible altered trajectory which would not leave the working area. The direction of movement bore no relation to the orientation of the rectangle. The alternate stimulus matched the initial stimulus except that one of the rectangles had either its orientation or its trajectory altered by ±90° (the ‘change type’ manipulation). Each stimulus was displayed for 700ms (a discussion of the precision of this timing is included below: **Error! Reference source not found.**). The alternate stimulus was either presented immediately after the initial stimulus’ 700ms display duration (‘unmasked’ condition) or after a 200ms mask (‘masked’ condition). During the mask the rectangles were rendered invisible, resulting in a plain white background. Crucially, all rectangles vanished and reappeared at the same time. Once the alternate stimulus had been displayed for 700ms the trial was restarted (Fig 1b), either immediately (unmasked condition) or following a second 200ms mask. Trials continued until the participant provided a response. A demonstration video showing 10 trials can be found at (<u>doi:10.6084/m9.figshare.5044894</u>).

**Fig 1.**
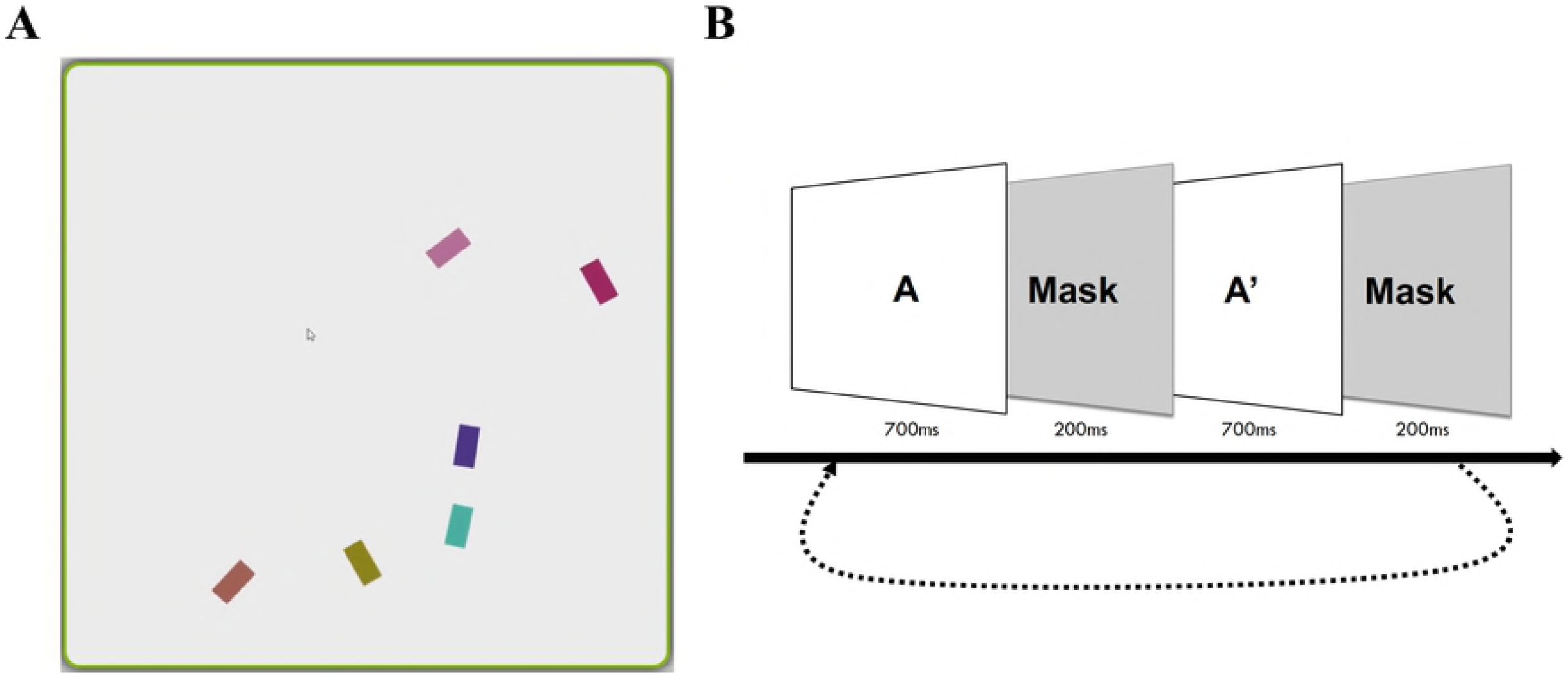
The task display and presentation procedure. a) Screenshot of the 700×700 pixel working area of the screen showing rectangles of random colours (selected from an approved region of colour-space) which moved in straight lines through the working area. After 700ms one of the rectangles would alter either its orientation or its trajectory by 90° either clockwise or anticlockwise. The task was to identify which of the rectangles had undergone this change. In the low load condition only two rectangles were presented; the high load condition presented six rectangles as shown. b) Flicker paradigm procedure. The initial scene (A) was displayed for 700ms, then a mask was put up for 200ms (masked condition) or 0ms (unmasked condition) before the initial scene was replaced with the altered scene (A’). The altered scene was displayed for 700ms and then masked and reverted to the initial scene. The process was repeated until the participant generated a response.

Research on multiple object tracking (28) has shown that, although object tracking does not interact with memory processes (29) (though see (30) for a dissenting view), tracked objects are to some extent resistant to change blindness (31,32).;tracking target selection is typically exogenous and based on colour and spatial location (33), As both of these were randomised in all trials, there was no systematic relationship between the likelihood of an object being tracked and its being the target object for that trial. Therefore, the dynamic paradigm did not undermine the validity of the change blindness.

Participants were asked to press the spacebar as soon as they had identified the altered rectangle. Pressing spacebar halted the movement of the rectangles and correct identification was checked by requiring the participants to click the rectangle which had changed. Participants were given the opportunity to practice the task until they were satisfied with their performance, and were provided with feedback as to their accuracy during the practice.

Each trial had one of eight possible types, defined by its specific arrangement of three different binary variables: whether a mask was present or not; whether scene load was low (2 rectangles) or high (6 rectangles); and whether the target rectangle was changed in orientation or trajectory. Experimental condition was selected randomly at the beginning of each trial. The probability of low scene load was 30%, chosen because more errors were expected under high load. Mask presence (present or absent) and change type (orientation or trajectory change) were equally likely for both options (50%). The outcome measure was time elapsed between the beginning of the first presentation of the altered state and the moment the spacebar was depressed.

After every 10 trials participants were provided with statistics showing their accuracy and average speed over the last 10 trials, as well as their averages for all trials thus far completed. This provided a sense of progress for participants, and encouraged them to focus on fast and accurate responses. At the end of all 50 trials participants were shown their average accuracy and speed, as well as the average accuracy and speed for all participants combined.

Participants were invited to complete as many trials as they wished, though the provision of full feedback after 50 trials was intended to incentivise the completion of at least 50 trials per participant. The overall number of trials, and the number of each type of trial seen by each participant was subject to some variation since some participants completed more trials than others and trial type was selected randomly at the beginning of each trial.

The lab participants completed 100 trials, and were given the information about their performance relative to others only after they have completed these 100 trials.

The task application was coded in HTML and JavaScript with the aid of the CraftyJS JavaScript game engine library version 0.7.0 (34). Results were sent using AJAX (Asynchronous JavaScript And XML) to a PHP script which stored them in a MySQL database and returned statistics to the participant when required. The JavaScript application was checked for compatibility with and parity between recent versions of the most common browsers (Microsoft Internet Explorer, Mozilla Firefox, Apple’s Safari, and Google Chrome). Statistical analysis of data was performed using R (35) and its tidyverse package (36). Initial analyses, reported in supplementary material, were performed using IBM SPSS Statistics package (version 22.0.0).

### Variation within the online cohort

The use of web-based psychological experiments is an increasingly popular approach to acquiring data which offers a number of trade-offs compared to laboratory experiments (37–39). Those advantages which are most salient to this project are the savings in time, money, and equipment, as well as the added convenience for participants who would have otherwise had to attend a laboratory session. Relevant disadvantages are primarily the result of non-standardised equipment (including screen brightness, size, and resolution) and environments (including noise levels and distractions) resulting in a slightly different experience of the experiment for each participant.

The paradigm was implemented with JavaScript. JavaScript’s control (used for stimulus timings) is less precise than other frequently-used languages (40). There was thus unsystematic variation in stimulus and mask durations of ±10ms, as well as some variation in the magnitude of this variation between browsers (on the order of ±5ms). The unsystematic variation constitutes noise, a factor addressed in the sample size (Participants). Variation on the basis of browser is handled statistically as a component of inter-participant variation (**Error! Reference source not found.**).

The retinal speed of stimuli was subject to variation between participants on the basis of screen size and viewing distance (which were not controlled). This variation is handled statistically as inter-participant variation (**Error! Reference source not found.**). A more pressing concern is the possibility that the experimental conditions may have been differentially affected by differences in stimulus retinal speed given that one of the conditions (trajectory change) was implemented through a change in the motion of the stimulus. This concern would be apt for manipulations in which the speed of the stimulus was altered (e.g. examining change detection for acceleration), but does not apply when, as here, the speed is kept constant while the direction of motion is altered.

The trade-off between these various factors was considered acceptable given the robustness of change blindness as a phenomenon: change blindness is inducible in a wide variety of situations from strict laboratory (31) to naturalistic (1) and virtual (41) settings. As expected given this robustness, pilot testing indicated that the experimental manipulations were successful in a range of hardware and software environments and usage scenarios. Furthermore, the online study was followed up by a lab-based replication using the same software in controlled conditions.

For the lab cohort, the above sources of variation were eliminated or reduced substantially (with the exception of the unsystematic variation in stimulus and mask durations caused by JavaScript’s imprecise timing control). In the lab-based part of the study the task was presented on a 37.1 cm x 33.3 cm inch Dell monitor with a resolution of 1280 x 1024, a refresh rate of 60Hz, and colour depth of 32 Bit. The participants completed the task seated at a comfortable distance from the monitor of approximately 57 centimetres.

### Paradigm selection

The flicker paradigm was selected because of its ease of implementation and its greater resilience to variations in screen size. This resilience is due to the particular method of transient suppression deployed in the flicker paradigm: in a flicker paradigm there are no change-specific transients since the transition (in the masked condition) is from initial stimulus, to mask, to alternate stimulus. Other methods, such as the mudsplash paradigm (12), rely upon attracting attention away from change-specific transients by introducing more salient transients elsewhere. While these other methods are likely to work given the robustness of change blindness as a phenomenon, screen size variation among online participants means that the distance between any two points on the screen could not be guaranteed to produce equal visual space distances for all participants. Thus, relying upon location-based transient-suppression would introduce unnecessary variation between participants.

## Results

### Descriptives and exclusions

Experimental data consisted of 3035 trials (online = 1545, lab = 1490) from 42 participants (online = 27, lab = 15). Trials were excluded if the wrong object was identified as having changed, if the trial took longer than 20s, or if the trial belonged to a participant who either had zero valid trials for any of the eight trial types or had an overall accuracy below 90% (Table 1).

**Table 1.** Trial exclusion process

| Criterion<br>(sequentially applied) | Trial count |  |  |  |
| --- | --- | --- | --- | --- |
|  | Online | Lab | Total | Running total |
| All trials | 2226 | 1746 | 3972 | **3972** |
| Errors | 120 | 109 | 229 | **3743** |
| RT > 20s | 139 | 81 | 220 | **3523** |
| Missing trial types | 190 | 0 | 190 | **3333** |
| Accuracy < 90% | 232 | 66 | 298 | **3035** |

The excluded trials were examined for differences in manipulation using χ^2^ tests with Yates’ continuity correction, conducted on those trials performed by the 42 participants included in the experimental data. Errors occurred significantly more often than expected by chance in masked than unmasked trials (χ^2^(1, *N* = 3354) = 30.0, *p* < .001), in high than low load trials (X^2^(1, *N* = 3354) = 29.5, *p* < .001), and in orientation than trajectory trials (χ^2^(1, *N* = 3354) = 5.67, *p* = .017).

Online participants completed an average of 57.2 (±SD 4.49) trials, and lab participants 99.3 (±SD 5.83). There was no significant difference in the number of orientation and trajectory trials included in the final statistical analysis for either the online (*t*(214.0) = −0.227, *M_diff_* = - 0.139 [95%CI: −1.35, 1.07], *p* = .821) or lab cohort (*t*(118.0) = −0.624, *M_diff_* = −0.667 [95%CI: - 2.78, 1.45], *p* = .534). The distribution of trials by contingency for each cohort is shown in **Error! Reference source not found.** These data, including excluded trials and participants, have been made available along with the script used to analyse them (https://doi.org/10.6084/m9.figshare.6580223).

**Fig 2.**
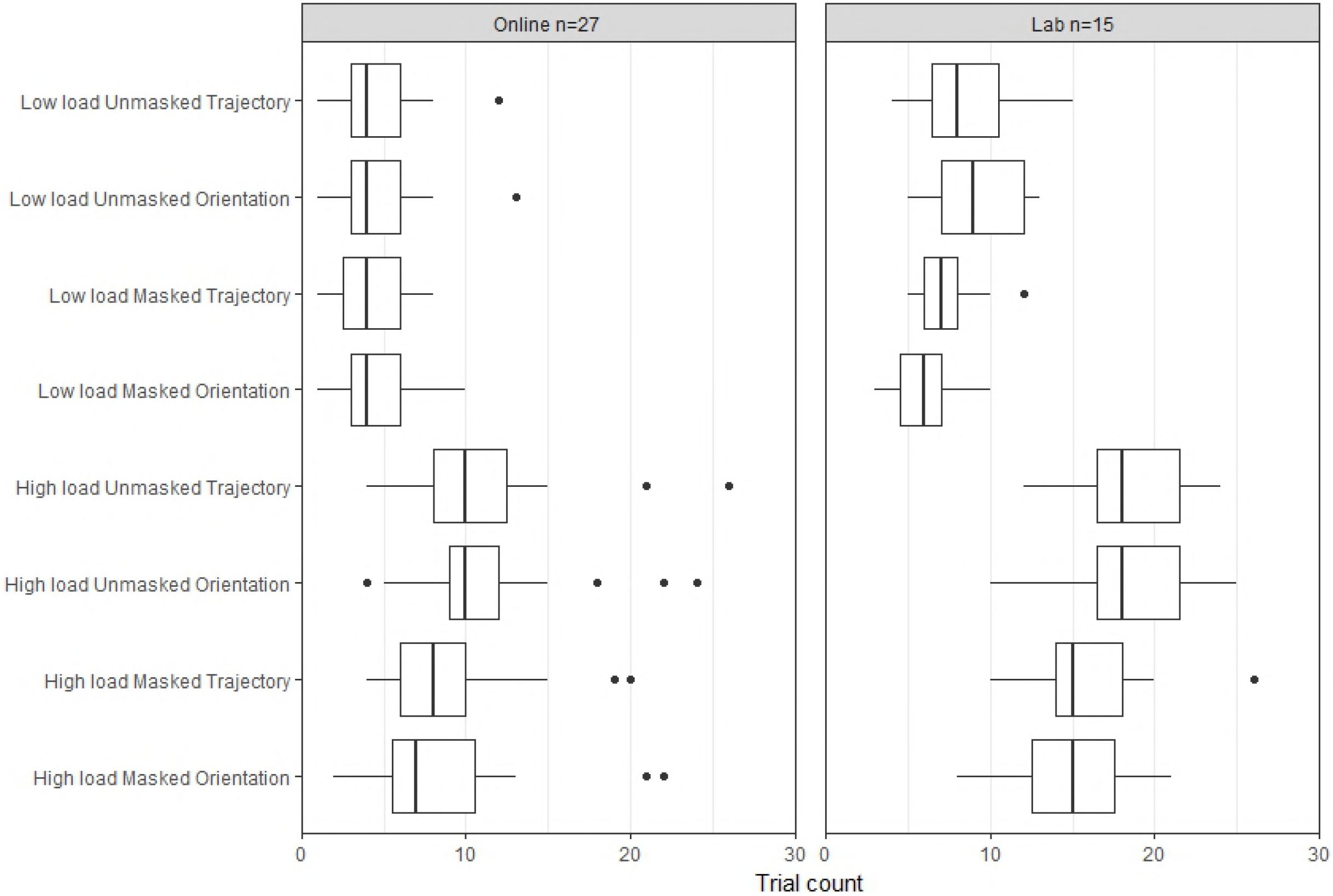
Trial type distribution. The trial type was selected at random at the beginning of each trial. Boxplots show the distribution of the number of trials of each trial type completed (successfully) by participants in the online and lab cohorts.

Analyses were performed with trials collapsed by participant. The means number of trials and response time for each condition, and their 95% confidence intervals, are shown in Table 2. Response times are calculated from the moment responding is enabled (after the altered stimulus becomes visible) until a response is recorded. For masked trials, in which repetitions of the mask make responding more difficult, response times are reduced by the duration of the masks displayed. Analysis using unadjusted response times made no difference to the pattern of results obtained.

**Table 2.** Means

| Trial Type |  | Number of Trials |  | Response Time (ms) |  |
| --- | --- | --- | --- | --- | --- |
|  |  | mean | 95%CI | mean | 95%CI |
| High load Unmasked Orientation | Online | 11.2 | 9.4, 12.9 | 1738.0 | 1105.9, 2370.1 |
|  | Lab | 18.5 | 16.4, 20.6 | 1574.4 | 1195.4, 1953.4 |
| Low load Masked Trajectory | Online | 4.3 | 3.5, 5.2 | 889.0 | 577.9, 1200.1 |
|  | Lab | 7.5 | 6.4, 8.5 | 762.9 | 637.1, 888.8 |
| Low load Unmasked Orientation | Online | 4.6 | 3.6, 5.6 | 813.3 | 638.4, 988.1 |
|  | Lab | 9.5 | 8.0, 10.9 | 722.4 | 604.8, 840.1 |
| High load Masked Orientation | Online | 8.1 | 6.3, 9.9 | 3786.4 | 3012.1, 4560.6 |
|  | Lab | 14.4 | 12.1, 16.7 | 4224.1 | 3597.0, 4851.3 |
| High load Unmasked Trajectory | Online | 11.0 | 9.2, 12.8 | 1282.7 | 942.7, 1622.7 |
|  | Lab | 18.6 | 16.7, 20.5 | 754.0 | 651.9, 856.0 |
| Low load Masked Orientation | Online | 4.5 | 3.5, 5.4 | 1207.4 | 735.9, 1678.9 |
|  | Lab | 5.9 | 4.8, 7.0 | 1034.7 | 814.4, 1255.0 |
| Low load Unmasked Trajectory | Online | 4.6 | 3.6, 5.6 | 655.3 | 498.7, 811.9 |
|  | Lab | 8.5 | 6.6, 10.5 | 486.7 | 449.6, 523.8 |
| High load Masked Trajectory | Online | 9.0 | 7.3, 10.6 | 3699.7 | 3002.1, 4397.3 |
|  | Lab | 16.4 | 14.0, 18.8 | 3565.5 | 3077.4, 4053.6 |

### Open science

Analysis of data from the lab cohort was preregistered (https://aspredicted.org/rh3d2.pdf) to use a 2×2×2 within-subjects ANOVA, and to be repeated covering both raw and adjusted response time as a dependant variable. These analyses were conducted, but, given the similarity to the main analysis reported below, are not reported in detail here. Detailed results from all analyses conducted in this paper, the raw data upon which they are based, and the scripts used to produce them, are available online https://doi.org/10.6084/m9.figshare.6580223.

### Main analysis

The data were analysed with a mixed (2×2×2×2) ANOVA. Change type (orientation vs trajectory), masking (unmasked vs masked), and load (low vs high) were the within-subjects variables, and cohort (online vs lab) was the between-subjects variable. Main effects were observed for all three within-subjects factors: responses were slower when masked (*F*(1,40) = 218.0, *p* < .001, η_p_^2^ = .846, *M_diff_* = −1358.7 [95%CI: −1677.7, −1039.7]); under high load (*F*(1,40) = 252.5, *p* < .001, η_p_^2^ = .863, *M_diff_* = −1750.6 [95%CI: −2046.7, −1454.4]); and for orientation changes (*F*(1,40) = 14.3, *p* < .001, η_p_^2^ = .285, *M_diff_* = 341.0 [95%CI: −689.6, 7.5]). A significant interaction was observed for masking x load, with the increase in response time for masked trials being exacerbated under high load (*F*(1,40) = 179.1, *p* < .001, η_p_^2^ = .820), while other interactions were not significant (all *F*(1,40) < 1.66, all *p* > .206, η_p_^2^ < .085). There was no effect of the between subjects factor, either as a main effect (*F*(1,40) = 0.232, *p* = .633, η_p_^2^ = .006, *M_diff_* = 118.4 [95%CI: −231.6, 468.3]), or in interactions (all *F*(1,40) < 2.50, all *p* > .122, all η_p_^2^ < .059).

The presence of an interaction between masking and load (Fig 3) indicates that change blindness occurred. The absence of three-way interaction between that interaction and change type is consistent with the suggestion that the change blindness effect is equivalent between change types. These data are consistent with the hypothesis that trajectory changes are detected and missed in a similar manner to orientation changes.

**Fig 3.**
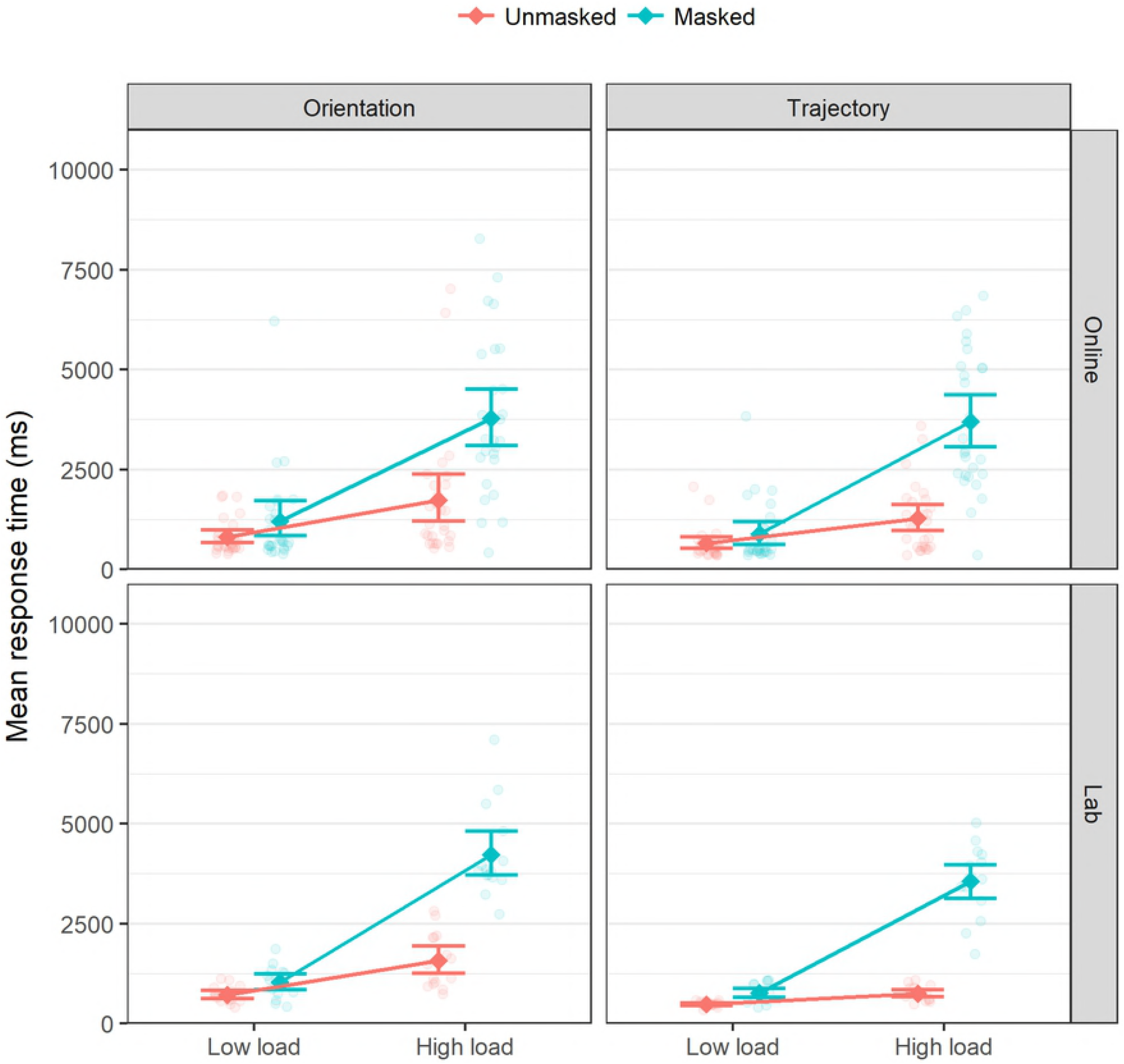
Change blindness interaction. The key pattern of interaction, whereby the response time for high load masked trials is far greater than either masked or high load trials alone, is evident for both orientation and trajectory changes. This relationship is stable between the online and lab cohorts. Error bars show 95% confidence intervals.

This analysis was checked for robustness under various different assumptions: using non-adjusted response times; excluding trials with a response time under 200ms (*N* = 18); including all trials with a non-erroneous response time; and analysing the cohorts separately. None of these alternate analyses resulted in a different core pattern of results (main effects of masking and load, and an interaction between them), though the main effect of change type was non-significant in some cases. For the lab cohort alone, a significant interaction arose between change type and load (η_p_^2^ = .414).

Finally, a reviewer noted that the dramatic disruption arising from refreshing the display after the A’ panel has finished (Fig 1b) constituted a larger disruption than the visual mask, and that results may be artificially elongated by this effect. Since this only occurs on trials where answers are not given within the first A’ panel (i.e. RT <= 700ms), we also performed analysis in which response time was replaced as the dependant variable by a binary variable indicating whether or not a response was made within the first A’ panel. The mean is thus the proportion of trial which were solved immediately, and is conceptually similar to measuring whether or not the participant detected the change in a one-shot paradigm.

The core result was robust to this analysis: unmasked changes were more likely to be noticed immediately than masked changes (*F*(1,40) = 200.8, *p* < .001, η_p_^2^ = .842, *M_diff_* = .302 [95%CI: .228, .377]); changes were more likely to be noticed immediately under low load (*F*(1,40) = 195.5, *p* < .001, η_p_^2^ = .830, *M_diff_* = .361 [95%CI: .289, .432]); and these effects interacted with one another (*F*(1,40) = 28.1, *p* < .001, η_p_^2^ = .414). Additionally, a main effect of change type was observed, with trajectory changes being noticed immediately more frequently than orientation changes (*F*(1,40) = 63.9, *p* < .001, η_p_^2^ = .670, *M_diff_* = .134 [95%CI: .054, .213]). This effect of change type also interacted with the masking x load interaction, resulting in a three way interaction (*F*(1,40) = 9.58, *p* = .004, η_p_^2^ = .229). The main effects of masking and change type interacted with cohort, both being increased by lab conditions (masking: *F*(1,40) = 11.7, *p* = .001, η_p_^2^ = .226; change type: *F*(1,40) = 17.3, *p* < .001, η_p_^2^ = .302).

## Discussion

The results demonstrate that change blindness was achieved using the implementation. Where scenes were sparsely populated enough for the relevant properties of all objects to be maintained in working memory, or where transients accompanying key changes were available, changes were noticed rapidly. Where working memory exhaustion and transient masking occurred simultaneously, changes were noticed far more slowly. This pattern of results is typical of change blindness in both direction and magnitude (2,13,27,42).

The use of dynamic stimuli did not compromise the orientation change blindness effects, consistent with suggestions of (26,43). The replication of change blindness to orientation changes validates the methodology used here, and the similar patterns of results in the orientation and trajectory conditions suggests that change blindness was also evoked for trajectory changes. The existence of change blindness to trajectory changes implies that trajectory changes are capable of directing attention exogenously since change blindness involves a loss of exogenous attention manipulation when its triggers are suppressed.

### Investigation of trajectory change blindness

Transients are not required to detect all changes, as occasional success in masked change detection tasks proves (44), and thus there is a question as to whether there are classes of short-term changes which are routinely detected without recourse to transients. The present study examines whether trajectory changes are a class of discriminable changes which do not depend upon the detection of transients or differences in patterns of transients.

Were trajectory change detection typically transient-dependant it would be expected that trajectory change detection response time would be modified in a similar way to orientation change detection response time under change blindness manipulations. This is demonstrated statistically by the absence of an interaction between the type of change variable and the load and masking variables. While certain choices in the statistical analysis did indicate differences in the magnitude of the change blindness interaction between orientation and trajectory changes, the larger effects were for the trajectory trials and the interaction remained strong for both change types. Since trajectory change detection responded in the same way as orientation change detection, and since orientation change detection is driven by the detection of transients whose presence the experiment directly manipulated, the conclusion is invited that trajectory change detection relies on the detection of visual transients (or patterns of transients) in the same way that orientation change detection does.

Orientation and direction-of-motion are neurophysiologically related under some conditions since the direction of slow movement is computed by direction-of-motion cells and rapid movement is computed by a combination of direction-of-motion and orientation-sensitive cells (45,46). Rapid motion was not a focus in the current design because the stimuli moved too slowly to produce motion streaks, but it is plausible that trajectory changes would be detected by the orientation of motion streaks in a rapid motion version of the current paradigm. In the study presented here it is likely that orientation and trajectory changes were detected by different neural populations: direction-of-motion cells for trajectory changes and orientation-sensitive cells for orientation changes.

The trajectory-orientation differences were stable across other manipulations, but were pronounced enough to produce a significant main effect of change type under some statistical choices. The difference may be explained by the consequences of the changes. Orientation changes happen ‘all at once’ in the sense that the orientation after the change is as different from the pre-change orientation as it will get. Trajectory changes are instantiated just as quickly when the direction of motion is altered, but the spatial position of the object differs increasingly from the extrapolated location along the original trajectory as time goes on. For changes which are not noticed immediately (due to transients orienting attention to the change), comparisons between incoming sensory data and an estimate derived by extrapolating from memory become more extreme over time for trajectory changes. This could result in trajectory changes being detected more quickly on average because the change is, in some sense, continuing to occur.

It is important to note here that the saliency of the trajectories was controlled through the use of objects moving on a two-dimensional plane and through randomisation of both initial trajectories and direction of deflection. It would be expected that in an experiment where changes increased the saliency of trajectories (e.g. to move towards the participant), those changes would be detected rapidly even where the change was masked. This expectation would be in keeping with previous findings suggesting a link between saliency and change blindness attenuation across various different kinds of saliency measures (2,18,19).

## Conclusion

This experiment replicates change blindness using a dynamic version of the flicker paradigm and shows that a pattern of results typical of change blindness can be obtained for trajectory changes.

The experiment demonstrates that detection of trajectory changes can be subject to change blindness. Change blindness is theorised to occur on the basis of the detection of transients, and thus this experiment can be taken to show that trajectory change detection depends on the detection of the fluctuations in the patterns of transients accompanying trajectory change. While there may be discriminations which can only be made using top-down mechanisms, the presence of transients driving a trajectory change suggests that bottom-up processes can account for, at the least, some discriminations regarding changes in higher-order object properties.

The differences between detection speed for orientation and trajectory changes suggest that trajectory changes are detected more readily and are slightly more resistant to masking by flicker. This may be due to the additional temporal information that expected trajectories afford.

Alternatively, trajectories of the separate elements may be represented in a gist-like pattern of movement which boosts the salience of a single trajectory deviation, whereas the orientation alterations may require serial search for identification of change. Future research examining eye movements during the detection of orientation and trajectory changes could further our understanding of this difference.

## Acknowledgements

Thanks are due to Alex Martin for vital feedback during the development of the testing application, for assistance in locating participants, and for helpful comments on various drafts of this manuscript. This work would have remained unpublished but for the insightful comments and guidance from Paul Graham. Thanks also to Jenny Bosten for particularly helpful comments and critique. Finally, the authors are grateful for the valuable contribution of three reviewers, which resulted in a clearer and better manuscript.

## Author Contributions

MJ designed the experiment, developed the application, analysed the data, and wrote the manuscript, all under the supervision of RC. NA conducted the lab-based replication and contributed to the drafting of the manuscript.

